# Emended description of the genus *Henriciella*: morphological and phylogenetic analysis identifies two subgenera with different reproductive strategies

**DOI:** 10.64898/2026.09.02.748869

**Authors:** Samantha Colby, Alyssa Cosklo, Eduardo Diazgranados, Natalie Fuller, Ana Garippa, Hiba Muhammed, Lauren Wall, Grace Washney, Mark L. Watson, David T. Kysela, Amelia M. Randich

**Author notes:** **Corresponding author and email address**, Amelia M. Randich.

## Abstract

*Henriciella* is the only genus with reported non-prosthecate members in the family *Hyphomonadaceae*, a large group of dimorphic prosthecate bacteria. Using type strains, we reinvestigated growth in media shown to support prosthecate morphotypes in marine caulobacters. We report that Marine Broth (MB) and Peptone Yeast Extract (PYE)-derived media supported prosthecate populations for all *Henriciella* species. Surprisingly, we observed that *H. aquimarina*, *H. mobilis*, and *H. pelagia* reproduce by tip budding whereas *H. marina* and *H. liltoralis* reproduce by asymmetric binary fission. These two observed reproductive strategies resemble those characterized for the model prosthecate bacteria *Hyphomonas neptunium* and *Caulobacter crescentus*. Phylogenomic analysis indicates that *H. marina*, *H. litoralis*, *H. algicola*, and *H. barbarensis* form a monophyletic subgenus that reproduces by asymmetric binary fission. We propose genus and species emendations that correct the record on this fascinating genus.

**Data summary:** 16S rRNA sequence data are available on Figshare as AB1 (https://doi.org/10.6084/m9.figshare.33072434) and FASTA files (https://doi.org/10.6084/m9.figshare.32732334).

## 4. Introduction

The *Caulobacteraceae* and *Hyphomonadaceae* are sister families of ubiquitous dimorphic prosthecate bacteria found in various aqueous and terrestrial environments (1–3). Members of both families are aerobic, chemoheterotrophic oligotrophs with complex developmental cycles, asymmetrical cell patterning, and cellular appendages called prosthecae. The family *Hyphomonadaceae* was created in 2005 to contain marine caulobacters that once belonged to *Caulobacteraceae* (4) and has been under considerable revision as new species accumulate and genomic data makes more sophisticated analysis possible. While both families were once included in the order Caulobacterales, the marine genera have been broken out into two new orders with three different families to recognize the phylogenomic distinction of these groups: the order Hyphomonadales (family *Hyphomonadaceae*) and the order Maricaulales (families *Robiginitomaculaceae* and *Maricaulaceae*) (5). The order Caulobacterales now only contains *Caulobacteraceae*. For simplicity, we will refer to the orders Caulobacterales, Hyphomonadales, and Maricaulales as the “CHM superclade” in this paper. Under this new classification scheme, *Hyphomonadaceae* currently includes only the genera *Hyphomonas*, *Hirschia, Henriciella*, and *Ponticaulis*. *Henriciella* was established in 2009 (6) and is the only genus in the *Hyphomonadaceae* with non-prosthecate members (1).

Alphaproteobacteria exhibit a great diversity of morphologies and reproductive strategies (8). This is largely due to the dimorphic cell cycle that pervades much of this large class of Pseudomonadota (Proteobacteria). In dimorphic bacteria, reproduction yields two cells that are morphologically and physiologically distinct from one another (9). The CHM superclade, Rhodobacterales, Hyphomicrobiales, and Sphingomonadales are all hypothesized to share an ancestral dimorphic cell cycle in which a sessile, surface-attached mother cell gives rise to a smaller, often motile, free-living “swarmer” daughter cell (**Figure 1A**) (10). Regulators of the core cell cycle network are highly conserved between the CHM superclade and the Hyphomicrobiales order, with simpler regulatory circuits detectable in more distant alphaproteobacterial orders where they likely coordinate morphology, division, motility, and other features (11). In the CHM superclade, the dimorphic cell cycle coordinates the development of two distinct appendaged (“prosthecate”) morphologies that reproduce by either asymmetrical binary fission or tip budding (**Figure 1B**). Understanding the variation in and evolutionary relationship of these prosthecate morphologies and reproductive strategies is a major step in understanding fundamental principles of bacterial growth and division.

**Figure 1.**
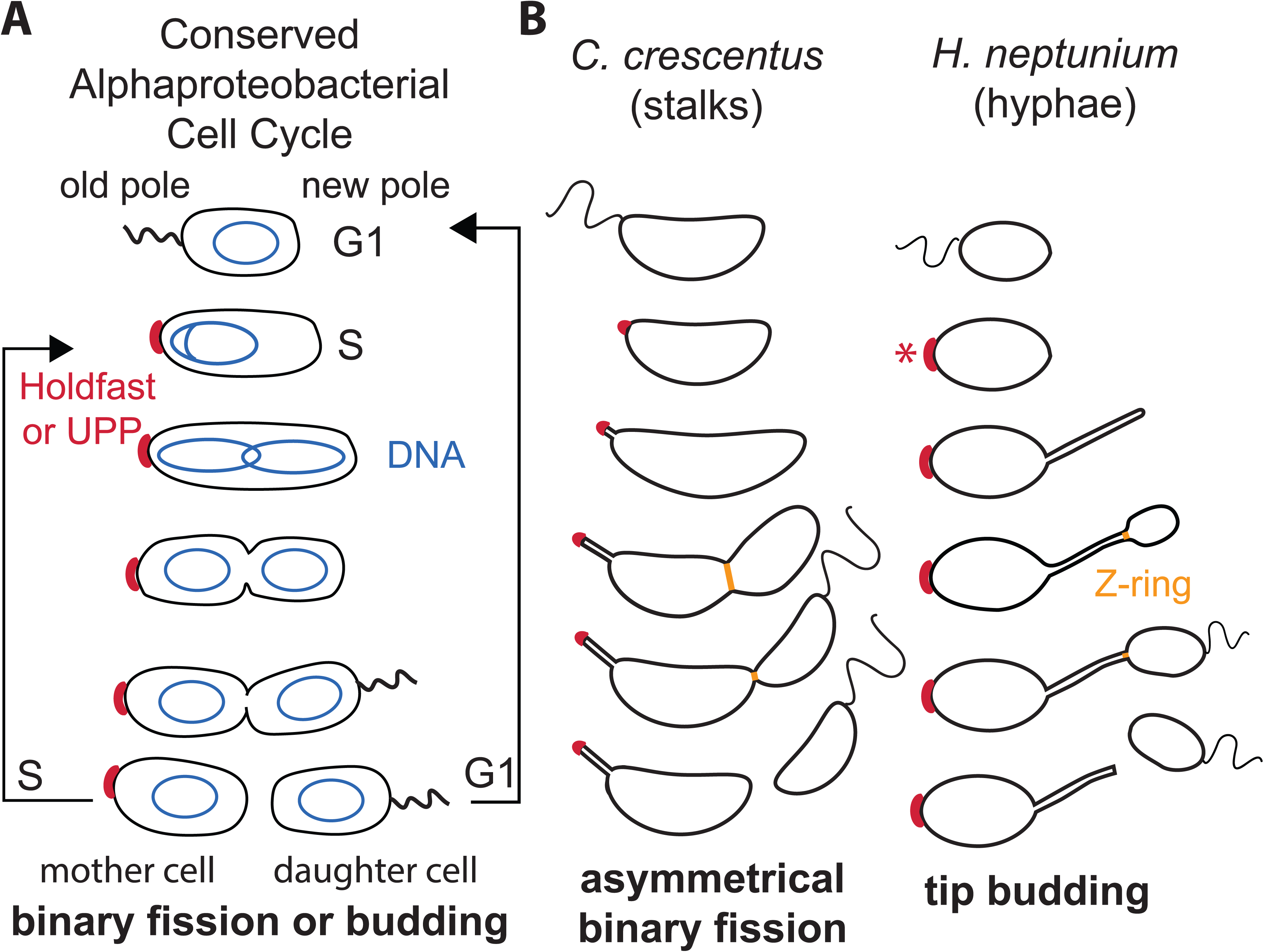
Asymmetrical binary fission and tip budding are built on the same ancestral alphaproteobacterial cell cycle. *(A)* Bacteria across the Caulobacterales, Rhodobacterales, Hyphomicrobiales (Rhizobiales), and Sphingomonadales orders of Alphaproteobacteria generally conserve an ancient biphasic cell cycle that, upon division, gives rise to a reproductive mother cell (S phase) and a motile, replication-inhibited daughter cell (G1 phase). This ancestral biphasic cell cycle is driven by asymmetrical polarity patterned by protein localization and phosphorylation gradients to define the old pole from the new pole. The new pole is designated as the pole produced with each division. After a specific time or due to environmental stimulation, the daughter cell will shift from G1 to S phase, start DNA (blue) replication, and produce an adhesin (red, “holdfast” in Caulobacterales, “UPP,” or unipolar polysaccharide, in Hyphomicrobiales, red) at the old pole. *(B)* The intersection of prosthecate morphologies with division gives rise to two different reproductive strategies in the Caulobacterales order, as represented by the model species *Caulobacter crescentus* and *Hyphomonas neptunium*. Although both species undergo dimorphic cell cycles, they produce prosthecae from different ends of the cell. This means that the prostheca of *C. crescentus* is produced at the old pole, is associated with the holdfast, and is therefore a “stalk.” These cells divide by asymmetrical binary fission where the z-ring forms near mid-cell (orange line). In contrast, the prostheca of *H. neptunium* is produced at the new pole, is associated with the daughter cell bud, and is therefore a “hypha.” These cells divide by tip budding, where the z-ring forms at the junction of the daughter bud and the hypha (orange line). Asterisk: *H. neptunium* does not produce a holdfast, but other *Hyphomonas* species do produce one at the old pole.

The word “prostheca” was coined by Staley in 1968 to distinguish cellular appendages that are narrow extensions of the entire cell envelope (inner membrane, cell wall, and outer membrane layers in Gram-negative bacteria) from other non-cellular appendages such as the slime stalks of *Nevskiaceae* or the fibrillar proteinaceous stalks of the Planctomycetes (12). This broad term covers a plethora of cellular appendages that appear throughout the bacterial domain, but especially in great variety in the Alphaproteobacteria (8). Within the CHM superclade, prosthecae are further distinguished by their relationship with the differentiated poles of the mother cell, and therefore their function: When produced at the old cell pole, prosthecae are associated with a sticky adhesin (holdfast) and called “stalks,” as exemplified by *Caulobacter crescentus* (**Figure 1B**). When produced at the new pole, they are associated with the generation of daughter buds and called “hyphae,” as exemplified by *Hyphomonas neptunium* (**Figure 1B**). Still further, prosthecae have been referred to as “pseudostalks” in *Asticcacaulis,* where they are not associated with either pole and are no longer adhesive or reproductive (13).

Although stalked caulobacters reproduce by asymmetrical binary fission and hyphate caulobacters reproduce by tip budding, they share the same ancestral cell cycle and polar patterning (**Figure 1**) (10,11). The evolutionary relationship of these two reproductive strategies is currently unknown, although tip budding is historically restricted to the orders Hyphomonadales and Maricaulales in the literature. The *Henriciella* genus (family *Hyphomonadaceae,* order Hyphomonadales) uniquely consists of a majority of reportedly non-prosthecate species dividing by binary fission. However, these species were characterized in Marine Broth (MB), the peptone levels of which have been shown to reduce the length and frequency of prosthecae in caulobacters (13). We sought to reassess the morphologies of the non-prosthecate *Henriciella* species in media known to support production of prosthecae and uniform dimorphic morphologies (3,14). We report here that all species are prosthecate, with some members reproducing by tip budding with hyphae and others reproducing by asymmetrical binary fission and producing stalks. These observations will help clarify the plasticity of these reproductive strategies.

## 5. Methods

### 5.1 Strains and Culture Conditions

Type species were obtained from the Japan Collection of Microorganisms (JCM), the Korean Collection of Type Cultures (KCTC), and the Belgian Coordinated Collections of Microorganisms (BCCM/LMG), or were generously provided by the Brun lab (**Table 1**). All strains were maintained as frozen 10% DMSO stocks and grown on Marine Broth (MB) or peptone yeast extract with sea salts (PYE+SS) plates with 1.5% agar at 26 or 30°C. Single colonies were used to inoculate liquid cultures. All species identities were confirmed by 16S sequencing (**Table 1**). Growth curves were constructed for each species in each medium and temperature from optical density measurements to determine exponential and stationary phase in each case. Bacteria were grown as 3-mL cultures at 21.5°C, 26°C, and 30°C shaking at 220 rpm.

**Table 1.** Verification of Strains by 16S Sequencing.

| Species | Strain | Type Strain | Acc. No | Identities | Gaps |
| --- | --- | --- | --- | --- | --- |
| *H. aquimarina* | LMG 24711* | Henriciella aquimarina strain P38 | NR_044573.1 | 1368/1370 (99.9%) | 2/1370 (0.1%) |
| *H. mobilis* | KCTC5 2576 | Henriciella mobilis strain M65 | NR_179031.1 | 1370/1372 (99.9%) | 2/1372 (0.1%) |
| *H. pelagia* | KCTC 52577 | Henriciella pelagia strain LA220 | NR_157792.1 | 1370/1370 (100%) | 0/1370 (0%) |
| *H. litoralis* | DSM 22014* | Henriciella litoralis strain SD10 | NR_116588.1 | 1326/1326 (100%) | 0/1326 (0%) |
| *H. marina* | JCM 15116 | Henriciella marina DSM 19595 strain Iso4 | NR_044345.1 | 1379/1382 (99.8%) | 3/1382 (0.2%) |
| *H. algicola* | LMG 29152 | Henriciella algicola strain MCS27 | NR_157788.1 | 1370/1371 (99.8%) | 0/1371 (0%) |
| *H. barbarensis* | LMG 28705 | Henriciella barbarensis strain MCS23 | NR_157787.1 | 1370/1371 (99.9%) | 1/1371 (0.1%) |

\*These strains were generously provided by the Brun Lab.

MB was purchased in the commercial formulation BD DIFCO 2216. Other media was formulated as indicated in **Table 2**. IOPYEM has been added to MediaDiv as https://mediadive.dsmz.de/medium/P11 (15). The enriched SPYEM medium (SPYEM+vm) was made by adding Vitamin Solution and Metals 44 1:1000 to SPYEM right before growth. For all media, tap water from Scranton, PA or Columbia, MO was purified by a Milli-Q Purification system. Instant Ocean sea salts (https://www.instantocean.com/) were used in place of Sigma Sea Salts due to the difference in cost; it had no effect or in some cases slight positive effects on cell morphology (data not shown). Optimized growth conditions for each species are listed in **Table 3**.

**Table 2.**
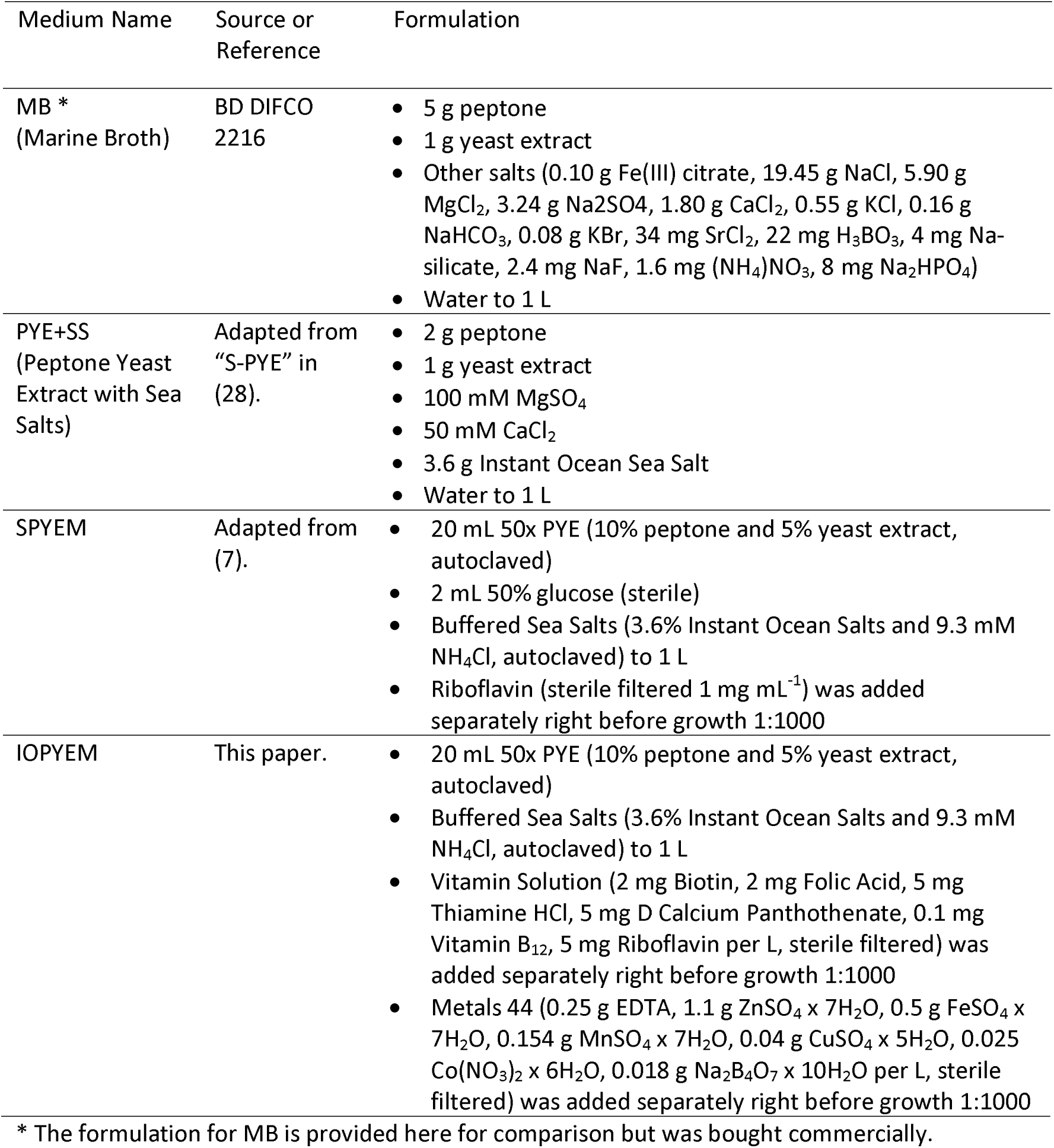
Media Composition.

**Table 3.** Quantification of Prosthecate Cells.

| Species | Optimized Growth Conditions | % Prosthecae (#cells/population)<sup>b</sup> | % With Secondary Prosthecae<sup>a</sup> (#cells/population)<sup>b</sup> |
| --- | --- | --- | --- |
| *H. aquimarina* | PYE+SS, 30°C | 39% (260/675) | Not seen in population |
| *H. mobilis* | PYE+SS, 30°C | 36% (270/752) | Exp<sup>c</sup> 0.1% (1/752), Stat 4.0% (37/919) |
| *H. pelagia* | PYE+SS, 30°C | 29% (240/830) | Exp<sup>c</sup> 0.8% (7/830), Stat 4.6% (37/800) |
| *H. litoralis* | IOPYEM, 21.5°C | 45% (346/761) | Not seen in population |
| *H. marina* | PYE+SS, 30°C | 22% (123/427) | Not seen in population |
| *H. algicola* | SPYEM, 30°C | 26% (206/785) | Not seen in population |
| *H. barbarensis* | SPYEM+vm, 21.5°C | 45% (397/879) | Not seen in population |

<sup>a</sup> “Secondary” refers to a prostheca that forms at the pole opposite of the primary prostheca.
<sup>b</sup> Counts were made from one biological replicate; however, prosthecae cells (and secondary prosthecae, where applicable) were always observed in cell populations across multiple independent experiments.
<sup>c</sup> “Exp” and “Stat” signify exponential phase and stationary phase.

### 5.2 16S Sequencing

All strains were grown on MA or PYE+SS plates at 30°C. Colony PCR was performed on single, well-isolated colonies using NEB OneTaq mix and IDT ReadyMade 16S sequencing primers “16S rRNA For” (AGA GTT TGA TCC TGG CTC AG) and “16S rRNA Rev” (ACG GCT ACC TTG TTA CGA CTT). Resulting amplicons were submitted to Plasmidsaurus for linear PCR sequencing (ONT). Sequences were confirmed by NCBI BLAST alignment and can be accessed at Figshare: https://figshare.com/s/aff01a622392f857af31.

### 5.3 Phase-Contrast Microscopy

Imaging was done using a Zeiss Axiophot upright microscope using Plan-NEOFLUAR 63X 1.25 NA oil Ph3 and Plan-NEOFLUAR 100X 1.3 NA oil Ph3 objectives outfitted with a Rigel monochrome camera with SONY back-illuminated CMOS. Cell in exponential and stationary phase were mounted on 1.5% (w/v) agarose pads made with sterile sea salts (3.6% Instant Ocean Salts, autoclaved). Cell measurements and quantification of cell morphotypes were done by hand using Fiji tools (16).

### 5.4 Transmission Electron Microscopy

Cells were grown to exponential phase before being pelleted at 7,000 xg and resuspended in fixative (2 % paraformaldehyde, 2 % glutaraldehyde in 100 mM sodium cacodylate buffer, pH 7.35). Unless otherwise stated, all reagents were purchased from Electron Microscopy Sciences and all specimen staining was performed at the Electron Microscopy Core Facility, University of Missouri. Fixed whole cells were washed by pelleting at 7,000 xg and resuspending in water three times. Samples were placed on negatively charged carbon coated copper grids. A negative charge was applied to the grid using a Pelco easiGlow Glow Discharge Cleaning System. Cells were allowed to settle on prepared grids for 2 minutes before removing the water with filter paper and replacing it with fresh water and allowing the cells to settle again for 2 minutes. Finally, water was replaced with 0.5% aqueous uranyl acetate for 30 seconds before drying and imaging. Images were acquired with a JEOL JEM 1400 transmission electron microscope (JEOL, Peabody, MA) at 80 kV on a Gatan Ultrascan 1000 CCD (Gatan, Inc, Pleasanton, CA) with the assistance of the University of Missouri EM Core.

### 5.5 Phylogenetic Analysis

NCBI reference genomes were downloaded as raw nucleotide assemblies. Phylosift (17) identified orthologs of 37 conserved protein coding genes, then aligned and concatenated the corresponding amino acid sequences. ModelTest-NG (18) selected the LG amino acid substitution model with gamma-distributed rate variation across sites and an additional invariant rate category, which was applied in all further analyses. MrBayes 3.2.7a (19) inferred the Bayesian consensus tree topology and clade posterior probabilities from two runs, each with four chains and heating parameter set to 0.05. Simulations spanned 300,000 generations, with 75,000 initial generations discarded for burn-in. Bootstrap support values were determined from 100 replicates using RAxML 8.2.9 (20), then mapped onto the Bayesian consensus topology.

## 6. Results and Discussion

Given that *H. algicola* and *H. barbarensis* were shown to produce stalks and divide by binary fission in a medium based on the *Caulobacter* medium PYE (7), we set out to verify whether the other species could also produce prosthecae in PYE-derived media. We characterized each species in both the medium in which it was first characterized, namely Marine Broth (MB, Difco 2216), and in PYE-derived media. Growth curves were determined for each species in each medium (data not shown) and morphology was assessed in exponential and stationary phase by phase-contrast microscopy. As described in the following sections, and shown in **Figures 2** and **3**, growth in MB resulted in morphologically heterogeneous populations with mother cells of various sizes and often with growth defects. MB contains 0.5% peptone (15), and these nutrient levels have been shown to cause morphological defects and reduction of the size or frequency of prosthecae in freshwater and marine caulobacters (3,13). In this work, PYE (0.2% peptone) made with sea salts (PYE+SS) and other PYE-derived media (SPYEM, IOPYEM, see Methods for formulations) supported more uniform growth and demonstrated that all *Henriciella* species produce prosthecae. However, more surprising was that some species divided by tip budding while others divided by binary fission.

**Figure 2.**
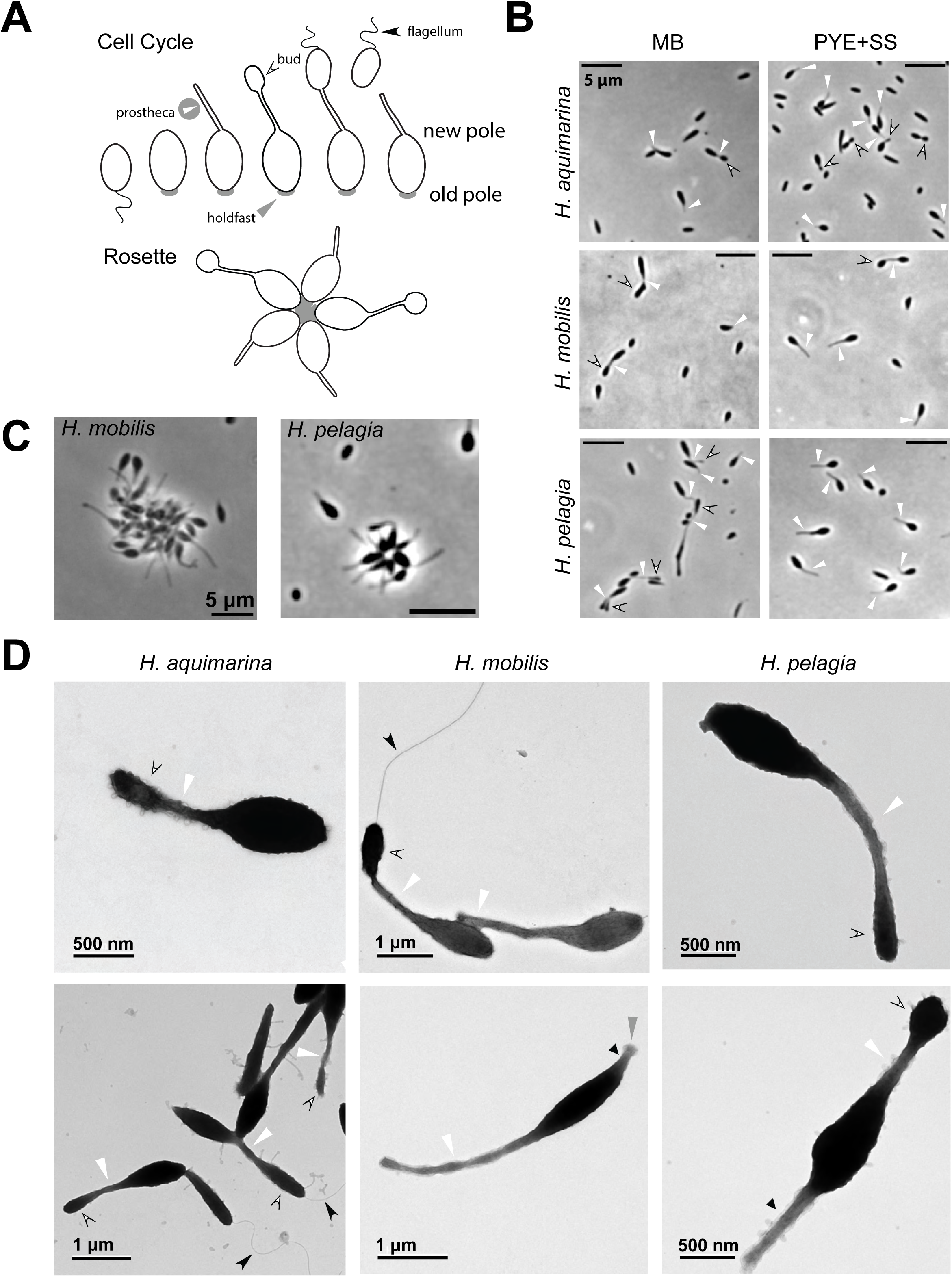
H. aquimarina, H. pelagia, and H. mobilis produce hyphae and reproduce by tip budding. (A) Schematic of the cell cycle and rosette formation for tip-budding Caulobacterales members. The holdfast appears in gray. *(B)* Phase-contrast images in Marine Broth (MB) and optimized medium (PYE+SS in all cases for these species). All phase-contrast scale bars are 5 µm. White arrowheads indicate prosthecae and black-outlined white, curved arrowheads indicate buds. *(C)* Phase-contrast images of rosettes formed by tip-budding species, with the prosthecae produced at the opposite pole to the holdfast. Scale bars are both 5 µm. *(D)* TEM images of *H. aquimarina*, *H. mobilis*, and *H. pelagia* grown in PYE+SS. Scale bars are as marked. Arrows are as in (B) with the addition of curved black arrowheads indicating flagella, short black arrowheads indicating secondary prosthecae, and gray arrowheads indicating holdfast.

**Figure 3.**
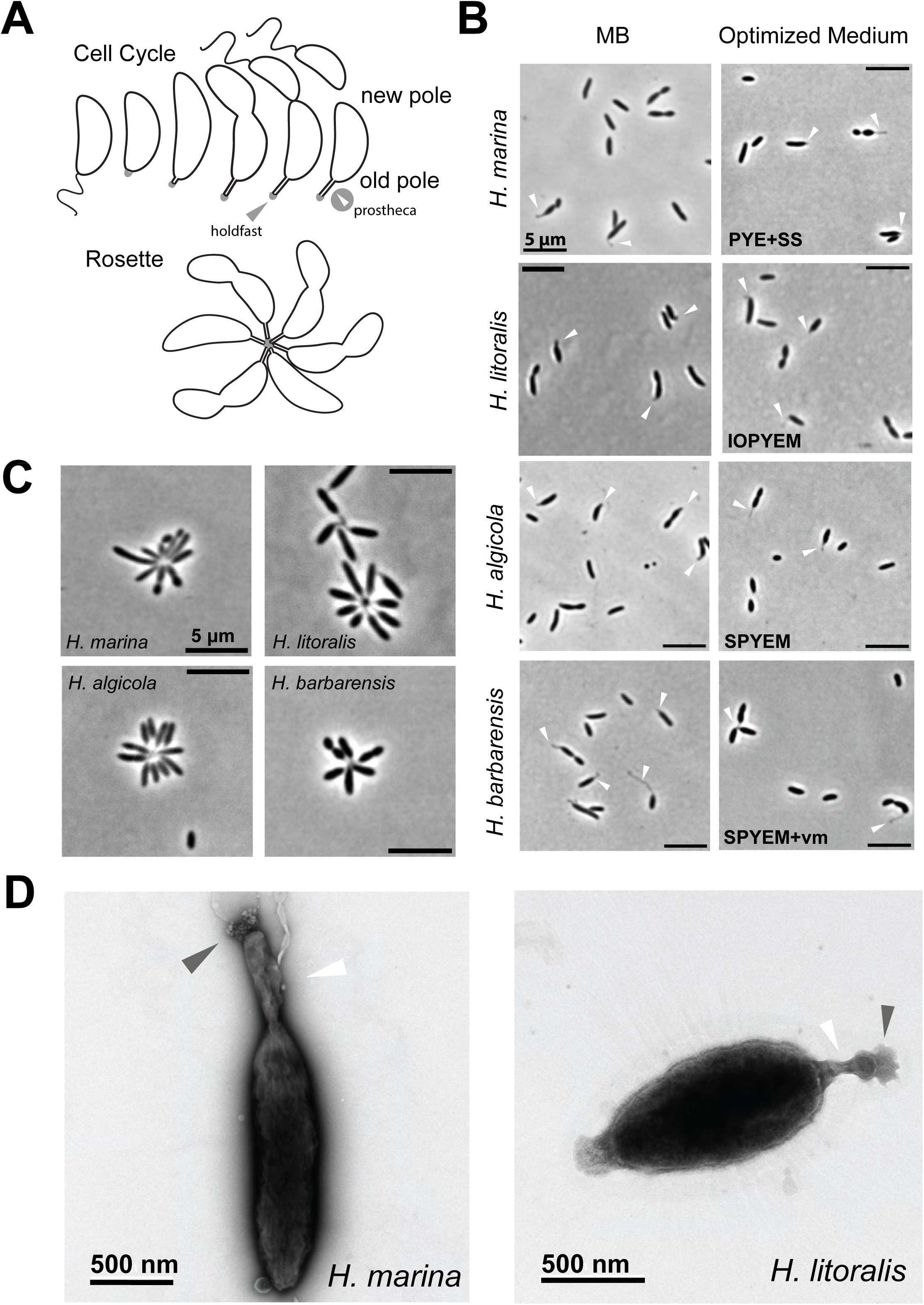
H. litoralis and H. marina produce stalks and reproduce by binary fission like H. barbarensis and H. algicola. *(A)* Schematic of the cell cycle and rosette formation for Caulobacterales members that divide by asymmetrical binary fission. The holdfast appears in gray. (B) Phase-contrast images in Marine Broth (MB) and optimal medium (PYE+SS, IOPYEM, SPYEM, or SPYEM+vm). All phase-contrast scale bars are 5 µm. White arrowheads indicate prosthecae. *(C)* Phase-contrast images of rosettes formed by these four species, with prosthecae produced at the same pole as the holdfast. *(D)* TEM images of *H. marina* grown in PYE+SS and *H. litoralis* grown in IOPYEM. Scale bars are 500 nm. White arrowheads indicate prosthecae and gray arrowheads indicate holdfasts.

### 6.1 H. aquimarina, H. pelagia, and H. mobilis produce hyphae and reproduce by tip budding

We observed that *H. aquimarina*, *H. pelagia*, and *H. mobilis* produced prosthecae and reproduced by tip budding. In tip-budding caulobacters, the prostheca (“hypha”) is associated with daughter buds and is opposite to the adhesive pole of the cell (**Figure 2A**). Typical tip-budding species can range from rod-shaped to teardrop-shaped and the most indicative morphotype of this reproductive strategy is the budding mother cell.

*H. aquimarina* was first identified as *Maribaculum marinum* by Lai *et al.* in 2009 and described as non-motile, non-budding, and non-prosthecate (21). It was moved to the *Henriciella* genus and renamed *H. aquimarina* by Lee *et al.* in 2011 (22). In MB, *H. aquimarina* grew as rods and teardrop-shaped cells that sometimes looked connected to smaller cells by prosthecae (**Figure 2B**). In wet regions of the agarose pad used as an imaging support, small swarmer cells could be seen swimming, indicating that *H. aquimarina* produces motile swarmer cells, like the rest of the CHM superclade. Although prosthecae were observable at the tapered end of the teardrop-shaped cells, buds were rare or difficult to discern (**Figure 2B**). Sometimes in exponential, but more so in stationary phase, the mother cells became elongated or had other developmental defects (data not shown for exponential phase, **Figure 2B**). Growth in PYE+SS did not change the shape of the growth curve (data not shown), but there were more unambiguously prosthecate cells and longer hyphae, which made buds easier to distinguish (**Figure 2B**). In PYE+SS in exponential phase, prosthecate (mother) cells were 1.3 ± 0.2 µm long and 0.64 ± 0.08 µm wide (N=20 cells, ± is standard deviation). 49% of cells had clearly identifiable prosthecae (**Table 3**). TEM confirmed that the appendages were extensions of the entire cell envelope with buds produced at their tips (**Figure 2D**). Flagella were often observed (lower image, **Figure 2D**), confirming a motile swarmer phase. No holdfasts were observed by TEM, in agreement with the notable absence of rosettes in both MB and PYE+SS cultures. However, large aggregates of tens to hundreds of randomly ordered cells formed in both media, suggesting that *H. aquimarina* likely produces a different type of EPS that is dispersed over the entirety of the cell body.

*H. mobilis* was initially described as growing as short motile rods (0.4-0.7 µm wide and 0.8-1.6 µm wide) with no mention of prosthecae or reproduction strategy (23). We observed prosthecae and reproduction by tip budding in MB, but the cells were often elongated and exhibited other morphological variability (**Figure 2B**). Similar to *H. aquimarina*, growth in PYE+SS did not alter the growth curve but greatly increased the uniformity of the mother cells, which were 2.1 ± 0.3 µm long and 0.87 ± 0.07 µm wide in exponential phase (N=20 cells, ± is standard deviation). 36% were prosthecate in these conditions (**Table 3**). TEM likewise confirmed that the appendages were prosthecae and produced buds (**Figure 2D**). Sometimes additional appendages were produced at the opposite end of the cell to the hyphae; 0.1% of exponential populations and 4% of stationary populations (**Table 3**). TEM verified that these were secondary prosthecae (3) and sometimes associated with a holdfast (**Figure 2D**). While holdfasts were not often seen by TEM, *H. mobilis* would occasionally make rosettes in PYE+SS but these often accumulated into larger aggregates (**Figure 2C**). This bacterium therefore may produce both a holdfast and another type of EPS all over the cell body such as that of *H. aquimarina*.

*H. pelagia* was also first described as short motile rods with no mention of prosthecae or reproduction strategy (24). Growth in MB produced prosthecate morphologies and clear signs of tip budding, but cells also exhibited severe developmental defects including extensive filamentation and ectopic poles (**Figure 2B**). Like the other tip-budding *Henriciella* species, its morphology became more uniform in PYE+SS where prosthecate (mother) cells were on average 1.8 ± 0.2 µm long and 0.7 ± 0.1 µm wide in exponential phase (N=20 cells, ± is standard deviation). Roughly 29% of its population was prosthecate in these conditions (**Table 3**). Rosettes were often observed in culture (**Figure 2C**) although holdfasts were not clearly identified in TEM. In these rosettes, hyphae are pointed outward, indicating that the holdfast is produced at the opposite pole of the cell, as expected for tip-budding caulobacters (25). Similar to *H. mobilis*, *H. pelagia* sometimes produced a secondary prostheca at the old pole that had similar proportions to the hypha (**Figure 2D**) and were present in 0.8% of exponential populations and 4.6% of stationary populations grown in PYE+SS (**Table 3**).

Collectively, our observations indicate that *H. aquimarina*, *H. mobilis*, and *H. pelagia* undergo the same cell cycle as *Hyphomonas*, the sister genus to *Henriciella* (**Figure 2A**). They all produce hyphae, which are extensions of the entire cell envelope that produce daughter buds. Two of the species, *H. mobilis* and *H. pelagia*, produced rosettes with prosthecae pointing outwards, suggesting that a holdfast is produced at the old pole, opposite to the hypha, in agreement with what is expected for this developmental cycle. Although the model tip-budding bacterium *Hyphomonas neptunium* does not produce a holdfast, other tip-budding relatives such as *Hyphomonas rosenbergii* (25) and *Hirschia baltica* (26) do, and the loss is considered recent (27). *H. mobilis* and *H. pelagia* also produced secondary prosthecae at the old pole, especially in stationary phase. *H. aquimarina* and *H. mobilis* produced higher-order aggregates, suggesting an additional adhesive EPS secreted all over the cell body, such as observed for *H. rosenbergii* (25). It is likely that all three species were morphologically mischaracterized due to their growth defects in MB. Previous researchers may possibly have based their observations for *H. mobilis* and *H. pelagia* on swarmer cells, which are generally more uniformly behaved in culture.

### 6.2 H. marina and H. litoralis produce stalks and reproduce by binary fission like H. algicola and H. barbarensis

We observed that *H. marina* and *H. litoralis* produced prosthecae and reproduced by asymmetrical binary fission like reported for *H. algicola* and *H. barbarensis*. In caulobacters that divide by asymmetrical binary fission and produce one polar prostheca, the prostheca (“stalk”) is associated with polar adhesin (“holdfast”) at the old pole (**Figure 3A**). Stalked species are generally rod-shaped with vibrioid and fusiform variants and the most indicative morphotype of this reproductive strategy is the pre-divisional, stalked mother cell.

*H. marina* was described as “motile, non-budding, non-stalked,” by Quan *et al.* in 2009 (6), but was described as having “hyphae” by Lee *et al.* in 2011 (22). The current species description also includes that “some cells form a mycelium, ranging from 7.8 to 8.0 µm in length” (6). We observed prosthecate cells in MB but found that growth in PYE+SS increased the number of observations (**Figure 3B**), with about 22% of the population producing stalks in exponential phase in PYE+SS (**Table 3**). In these conditions, prosthecate (mother) cells were 1.5 ± 0.3 µm long and 0.60 ± 0.08 µm wide (N=20 cells, ± is standard deviation). *H. marina* readily formed rosettes with the stalks oriented to the inside (**Figure 3C**), as expected for caulobacters with one polar stalk (**Figure 3A**). TEM imaging confirmed that the appendages were prosthecae and often associated with large holdfasts (**Figure 3D**). We never observed any “mycelia” in any medium.

*H. litoralis* was reported as motile rods that divide by binary fission (22). In MB, this species grew as rods with odd, tapered ends at one pole that could be distorted prosthecae (**Figure 3B**). Filamentation was common and some cells were so misshapen it was difficult to determine if the reproductive strategy was binary fission. Cells grown in PYE+SS were slightly curved or kinked and produced more noticeable appendages at one pole, but these were wider than expected for prosthecae (data not shown). We therefore tried SPYEM, which had been used successfully to grow *H. algicola* and *H. barbarensis* (7). This medium is formulated as PYE with buffered sea salts (9.3 mM NH_4_Cl), glucose (0.1%), and riboflavin (1 µg/mL). In SPYEM, the cells were not kinked but still produced wide, short appendages (data not shown). We formulated a similar buffered PYE medium called IOPYEM without glucose but with the addition of multiple vitamins and trace metals (see Methods). In this medium, the cells took on a shorter rod and liminoid shapes, produced thinner, more distinct prosthecae, and clearly divided by binary fission (**Figure 3B**). In IOPYEM in exponential phase, 45% of the population produced prosthecae (**Table 3**) and prosthecate (mother) cells were 2.1 ± 0.4 µm long and 0.76 ± 0.06 µm wide (N=20 cells, ± is standard deviation). This species readily produced rosettes (**Figure 3C**). Some of the cells appeared to produce secondary prosthecae at the old pole, but TEM imaging indicated that only one end produced a prostheca, which was often associated with a distinct holdfast (**Figure 3D**). The new pole sometimes had additional outer envelope material but was not an extension of the entire envelope (**Figure 3D**).

Previous characterization of *H. algicola* and *H.* barbarensis in SPYEM reported them to be prosthecate and to form rosettes using a holdfast at the end of a stalk (7). We grew these species in MB and PYE-derived media to assess their similarity to *H. marina* and *H. litoralis*. In MB, *H. algicola* also produced shorter, fatter stalks (**Figure 3B**). Moreover, both species spheroplasted in late exponential phase when grown in MB. Growth in SPYEM, the original medium for characterization by Abraham *et al.* (7) mostly alleviated both problems. In SPYEM in exponential phase, 26% of the *H. algicola* population was prosthecate (**Table 3**). Adding additional vitamins and trace metals to SPYEM (SPYEM+vm) and growing at room temperature (21.5°C) further promoted uniform cellular morphologies and stalk frequency for *H. barbarensis*. Under these conditions, 45% of the *H. barbarensis* population was prosthecate (**Table 3**). We confirmed that these species also produce rosettes with inward-pointing prosthecae (**Figure 3C**).

In sum, we found that *H. marina* and *H. litoralis* were morphologically very similar to *H. algicola* and *H. barbarensis* and that all four undergo the same cell cycle as *Caulobacter crescentus* and its marine relatives that divide by asymmetrical binary fission (**Figure 3A**). They all produced stalks, which are extensions of the entire cell envelope that are associated with the holdfast at the old pole, at similar frequencies as *H. algicola* and *H. barbarensis* (**Table 3**). Unlike the tip-budding *Henriciella* species, these stalked *Henriciella* species were more sensitive to media conditions and often responded well to the addition of vitamins and trace metals. Under these optimized growth conditions, we did not observe any tip budding. It is likely that *H. litoralis* and *H. marina* were previously mischaracterized as non-prosthecate due to assessment in MB.

### 6.3 The Henriciella species that divide by binary fission form a monophyletic subgenus

To evaluate how these two different reproductive strategies related to each other in the *Henriciella* genus, we mapped the reproductive strategy to a maximum likelihood tree constructed from concatenated gene sets for each species and other CHM superclade members (**Figure 4**). The branch order of this tree matches that of Ren *et al*. (23). In this tree, the stalked species *H. litoralis*, *H. marina*, *H. algicola*, and *H. barbarensis* form a monophyletic clade within the *Henriciella* genus, suggesting tip budding was lost in a common ancestor. The rest of the genus divides by tip budding like the sister genus *Hyphomonas* and the close relative *Hirschia*.

**Figure 4.**
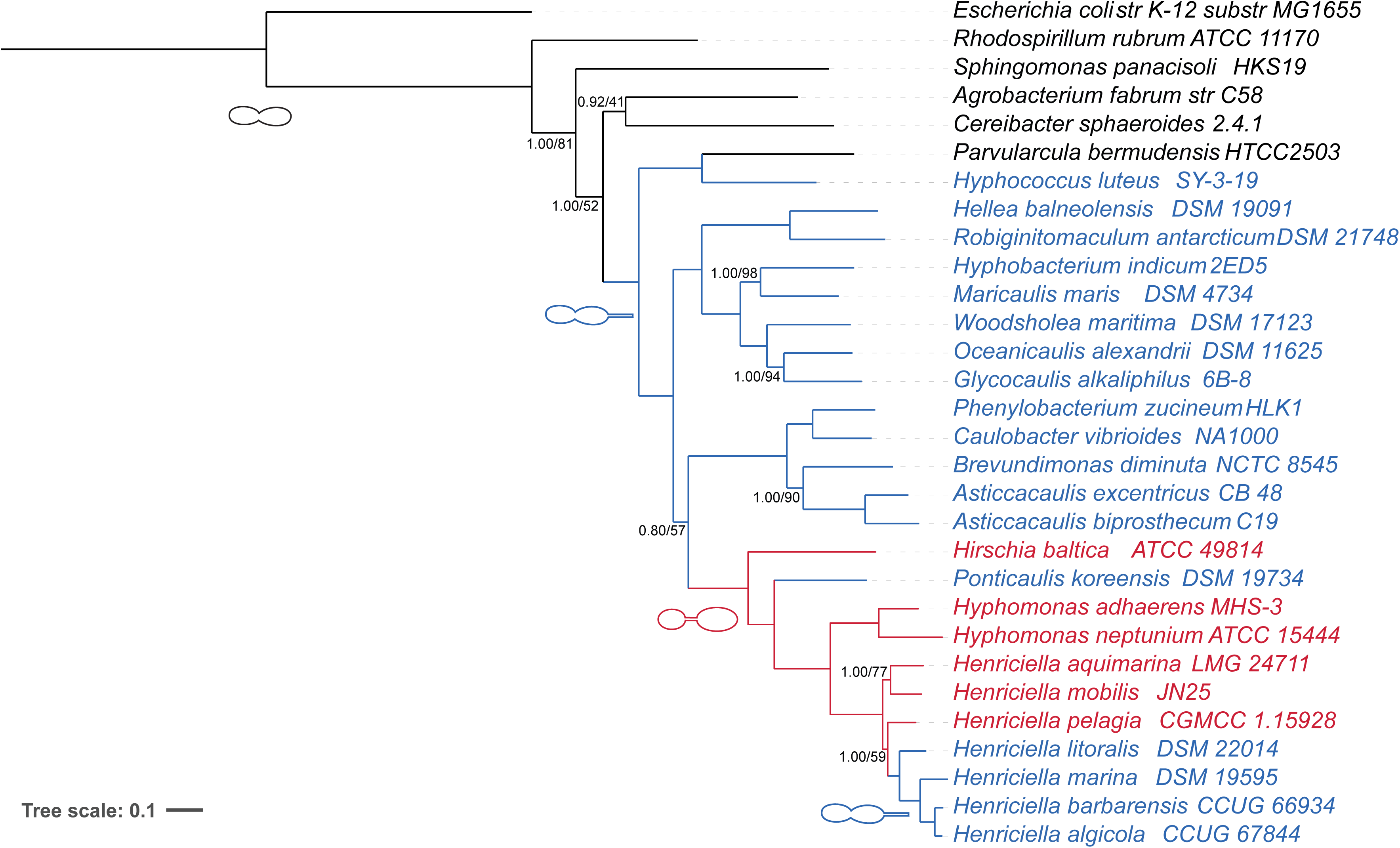
The Henriciella species that divide by binary fission form a monophyletic subgenus. Maximum likelihood tree derived from concatenated genome sequence data. Cell schematics and colors indicate inferred ancestral morphologies and their inheritance: black branches are non-prosthecate taxa, blue branches are taxa that reproduce by asymmetrical binary fission and produce stalks, red branches are taxa that reproduce by tip budding and produce hyphae. A concatenated alignment of 37 conserved protein sequences was used to infer the Bayesian consensus phylogeny shown. Support values are shown along branches as Bayesian posterior probability/bootstrap value. Only branches with posterior probability below 1.00 or bootstrap value below 100 are indicated; all remaining branches are fully supported by both metrics.

### 6.4 Concluding Remarks

Contrary to previous reports, we have determined that all current members of the genus *Henriciella* produce prosthecae. More noteworthy, we observed that three members reproduce by tip budding, like the sister genus *Hyphomonas*, and that four members form a monophyletic subgenus that reproduces by asymmetrical binary fission. It is likely that the prosthecate morphologies and the three cases of tip budding were missed on initial investigation due to characterization in Marine Broth, which, while standard for growth of marine bacteria, contains too much peptone for normal growth of these oligotrophic species.

We hope that our work serves as a guide for characterization of future CHM superclade isolates, of which there will be many more as metagenomic studies have already identified a great number of uncultured species that cluster with Hyphomonadales and Maricaulales members (2,10). The ecological niches of CHM superclade members are vast, including not only ocean and freshwater, but also soils, leaf litter, mineral deposits, and ectocommensal relationships with various other microorganisms (3,14). Any future isolates that genetically cluster with the CHM superclade should be assessed for morphological and reproductive traits in media with less than 0.5% soluble organic nutrients. In addition, attention should be paid to the likely dimorphic cell cycles of these new isolates, which often have a prosthecate mother cell and a flagellated daughter cell. Variation in and absence of these traits is of great interest to the communities that study the CHM superclade specifically, and bacterial growth, division, and cellular development in general.

Given our observations, we propose the following emendations:

## 7. Genus and Species Emendations

### 7.1 Emended Description of the Genus Henriciella Abraham et al. 2017

The entire emended description is given here. The description of this genus in terms of its developmental cycle, reproductive strategy and morphological characteristics is modified from that given by Quan *et al.* (6) and emended by Lee *et al.* (22) and Abraham *et al.* (7): Cells are Gram-negative, aerobic, and non-spore-forming with a dimorphic prosthecate cell cycle in which mother cells produce a prostheca and motile daughter cells possess one polar flagellum. Prosthecae vary in length depending on the species and environmental conditions and extend from one pole as a continuation of the long axis of the cell. However, some species reproduce by tip budding, where the daughter cell buds from the end of a prostheca (hypha) formed at the pole produced by division. A subset of these tip-budding species produce adhesive material (holdfast) at the opposite pole to the prostheca. Other species reproduce by asymmetrical binary fission and produce a prostheca (stalk) with adhesive material (holdfast) at its distal end. Colonies are circular, convex, and range from colourless to yellow. Chemoorganotrophic, aerobes, can grow anaerobically probably using amino acids as fermentable carbon sources. Cells can store carbon as poly-β-hydroxybutyrate and show activity of α-glucosidase. Species have complex growth requirements and grow on peptone-yeast extract media with 301g L^−1^ NaCl, with optimal growth between 10–50 g L^−1^ NaCl. Peptone levels at 0.5% and above cause morphological defects. No growth occurs with salt concentrations at or below 51g L^−1^. The temperature range for growth is 10–40 °C, 20–35 °C optimal. Polar lipids are α-d-glucopyranosyl diacylglycerol, α-d-glucopyranuronosyl diacylglycerol, phosphatidyl diacylglycerol and α-d-glucuronopyranosyl diacylglycerol taurineamide. The major fatty acid is C18 : 1ω7c; the major respiratory quinone is Q-10. The DNA G+C content is 55.2–62.0 mol%.

The type species is *Henriciella marina*.

### 7.2 Emended Description of the Species Henriciella aquimarina Lee et al. 2011

The entire emended description is given here. The description of this species in terms of its developmental cycle, reproductive strategy and morphological characteristics is modified from that given by Lai *et al.* (21) and emended by Lee *et al.* (22): Cells are Gram-negative, aerobic, non-spore-forming, short rods or teardrop-shaped cells with a dimorphic prosthecate cell cycle in which mother cells produce one hypha and motile daughter cells possess one polar flagellum. Cells reproduce by tip budding, where the daughter cell buds from the hypha of the mother cell. Holdfasts are not present. Carotenoid, bacteriochlorophyll a and the genes for anoxygenic photosynthesis (pufLM) are not found. Chemoheterotrophic. Cells are 1.2–1.4 μm long and 0.8–0.9 μm wide. Negative for lipase (Tween 80), amylase, urease, gelatinase, arginine dihydrolase and indole, hydrolysis of aesculin and reduction of nitrate to nitrite. On 216L agar, forms smooth, grey colonies with regular edges that are 2–3 mm in diameter after 72 h incubation at 28 °C. Grows with 2–12 % NaCl (optimum 3–10 %) and at 10–42 °C (optimum 25–37 °C), but not at 4 or 45 °C after 7 days. Unable to ferment glucose. The predominant fatty acids are C16 : 0, C17 : 0, C18 : 1 ω7c, C18 : 0 and C18 : 1 ω7c 11-methyl. Sensitive to (μg per disc unless otherwise stated) co-trimoxazole (25), ofloxacin (5), tetracycline (30), minomycin (30), penicillin G (10), streptomycin (10), ciprofloxacin (5), neomycin (10), vibramycin (30), piperacillin (100), rocephin (30), kanamycin (30), ampicillin (10), gentamicin (10), rifampicin (5), cephradin (30), chloromycetin (30), carbenicillin (100) and erythromycin (15). Resistant to cephalexin (30), cephazolin (30), cefobid (30), clindamycin (2), furazolidone (15), lincomycin (2), metronidazole (5), norfloxacin (10), oxacillin (1), polymyxin B (30 U) and vancomycin (30). With API ZYM, positive for acid phosphatase, alkaline phosphatase, cystine aminopeptidase, leucine aminopeptidase, naphthol-AS-BI-phosphoamidase, trypsin, valine aminopeptidase, α-chymotrypsin and α-glucosidase; weakly positive for esterase (C4), esterase lipase (C8) and lipase (C14); negative for N-acetyl-β-glucosaminidase, α-fucosidase, α- and β-galactosidase, α-mannosidase, β-glucosidase and β-glucuronidase. With API 20NE, does not utilize adipic acid, capric acid, d-glucose, maltose, d-mannitol, d-mannose, l-arabinose, malic acid, N-acetylglucosamine, phenylacetic acid, potassium gluconate or trisodium citrate. With GN2 MicroPlates, positive for utilization of l-aspartic acid, l-glutamic acid, l-ornithine, l-threonine, Tweens 40 and 80, α-ketobutyric acid, α-ketoglutaric acid, α-ketovaleric acid and β-hydroxybutyric acid; weakly positive for utilization of acetic acid, cis-aconitic acid, citric acid, d-alanine, d-arabitol, cellobiose, dextrin, d-galacturonic acid, d-serine, glucuronamide, glycogen, glycyl l-aspartic acid, glycyl l-glutamic acid, lactulose, l-alaninamide, l-alanine, l-alanyl glycine, l-asparagine, l-leucine, l-serine, maltose, pyruvic acid methyl ester, succinic acid monomethyl ester, phenylethylamine, propionic acid, quinic acid, α-d-glucose, γ-aminobutyric acid and γ-hydroxybutyric acid; negative for utilization of 2,3-butanediol, 2-aminoethanol, adonitol, bromosuccinic acid, dl-carnitine, dl-lactic acid, dl-α-glycerol phosphate, d-fructose, d-galactonic acid lactone, d-galactose, d-gluconic acid, d-glucosaminic acid, d-glucuronic acid, d-mannitol, d-mannose, melibiose, d-psicose, raffinose, d-saccharic acid, d-sorbitol, trehalose, formic acid, gentiobiose, α-d-glucose 1-phosphate, d-glucose 6-phosphate, glycerol, hydroxy-l-proline, i-erythritol, inosine, itaconic acid, l-arabinose, l-fucose, l-histidine, l-phenylalanine, l-proline, l-pyroglutamic acid, l-rhamnose, malonic acid, myo-inositol, N-acetyl-d-galactosamine, N-acetyl-d-glucosamine, p-hydroxyphenylacetic acid, putrescine, sebacic acid, succinamic acid, succinic acid, sucrose, thymidine, turanose, uridine, urocanic acid, xylitol, α-cyclodextrin, α-lactose, α-hydroxybutyric acid and methyl β-d-glucoside.

The type strain, P38^T^ (=CCTCC AB 208227^T^=LMG 24711^T^=MCCC 1A01086^T^), was isolated from deep seawater of the Indian Ocean.

### 7.3 Emended Description of the Species Henriciella litoralis Lee et al. 2011

As given by Lee *et al.* (22) with the following amendments: Cells are Gram-negative, strictly aerobic, liminoid to fusiform rods with a dimorphic prosthecate cell cycle in which mother cells produce a stalk and motile daughter cells possess one polar flagellum. Cells reproduce by asymmetrical binary fission and produce a stalk with adhesive material (a holdfast) at its distal end.

### 7.4 Emended Description of the Species Henriciella marina Quan et al. 2009

The entire emended description is given here. The description of this species in terms of its developmental cycle, reproductive strategy and morphological characteristics is modified from that given by Quan *et al.* (6): Cells are Gram-negative, strictly aerobic, short rods with a dimorphic prosthecate cell cycle in which mother cells produce a stalk and motile daughter cells possess one polar flagellum. Cells reproduce by asymmetrical binary fission and produce a stalk with adhesive material (a holdfast) at its distal end. Cells are usually 0.4∼0.7 µm wide and 0.7∼2.3 µm long. Good growth occurs on R2A with 1% NaCl, MA, and *Caulobacter* peptone yeast extract media with 3.6% sea salts. Colonies are translucent and shiny. Growth occurs at 10∼ 37°C (optimum, 20°C), at pH 5.3∼10.5 (optimum, pH 6.9∼ 7.6), and in the presence of 1∼15% NaCl (optimum, 1∼2%). In the API ZYM system, positive for alkaline phosphatase, esterase (C4), esterase lipase (C8), lipase (C14), leucine ary-lamidase, valine arylamidase, trypsin, α-chymotrypsin, adn acid phosphatase, α-glucosidase, α-fucosidase, and naphthol-AS-BI-phosphohydrolase activities; negative for cystine arylamidase, α-galactosidase, β-galactosidase, β-glucuronidase, β-glucosidase, and α-mannosidase *N*-acetyl-β-glucosaminidase activities. In the API 20NE system, negative for nitrate re-duction, indole production, glucose acidification, β-galacto-sidase, and arginine dihydrolase activities; do not hydrolyze aesculin, urea, and gelatin; do not use glucose, arabinose, mannose, mannitol, *N*-acetyl-glucosamine, maltose, gluconate, caprate, adipate, malate, citrate, and phenyl-acetate as a sole carbon source. The major cellular fatty acids are C18:1 ω7c (53.5%), C17:1 ω5c (11.7%), C17:1 ω6c (8.1%), C16:0 (7.8%), C17:0 (4.8%), C15:0 (2.9%), C16:1 ω5c (2.2%), summer feature 3 (1.7%), and C12:0 3-OH (1.1%).

The type strain Iso4^T^ (=KCTC 12513^T^ = DSM 19595^T^ = JCM 15116^T^) was isolated from seawater in the East Sea of Korea, at a depth of 100 m.

### 7.5 Emended Description of the Species Henriciella mobilis Ren et al. 2021

As given by Ren *et al.* (23) with the following amendments: Cells are Gram-negative, aerobic, non-spore-forming, short rods or teardrop-shaped cells with a dimorphic prosthecate cell cycle in which mother cells produce one hypha and motile daughter cells possess one polar flagellum. Cells reproduce by tip budding, where the daughter cell buds from the hypha of the mother cell. Holdfasts are present at the opposite end of the cell to the hypha and cells produce large aggregates. Some cells produce a secondary prostheca at the same pole as the holdfast.

### 7.6 Emended Description of the Species Henriciella pelagia Wu et al. 2017

As given by Wu *et al.* (24) with the following amendments: Cells are Gram-negative, aerobic, non-spore-forming, short rods or teardrop-shaped cells with a dimorphic prosthecate cell cycle in which mother cells produce one hypha and motile daughter cells possess one polar flagellum. Cells reproduce by tip budding, where the daughter cell buds from the hypha of the mother cell. Holdfasts are present at the opposite end of the cell to the hypha and cells produce rosettes. Some cells produce a secondary prostheca at the same pole as the holdfast.

## 8. Author statements

### 8.1 Author contributions

Samantha Colby: Formal analysis, Investigation. Alyssa Cosklo: Formal analysis, Investigation, Funding acquisition. Eduardo Diazgranados: Formal analysis, Investigation. Natalie Fuller: Formal analysis. Ana Garippa: Formal analysis, Investigation. Hiba Muhammed: Formal analysis, Investigation, Funding acquisition. Mark Watson: Investigation, Funding acquisition. Lauren Walls: Investigation. Grace Washney: Formal analysis, Investigation, Funding acquisition. David Kysela: Methodology, Software, Formal analysis, Investigation, Writing — review & editing. Amelia M. Randich: Conceptualization, Methodology, Validation, Formal analysis, Investigation, Resources, Data curation, Writing — original draft, Writing — review & editing, Visualization, Supervision, Project administration, Funding acquisition.

Role definitions:

- Conceptualization: https://casrai.org/credit/roles/conceptualization

- Data curation: https://casrai.org/credit/roles/data-curation

- Formal analysis: https://casrai.org/credit/roles/formal-analysis

- Funding acquisition: https://casrai.org/credit/roles/funding-acquisition

- Investigation: https://casrai.org/credit/roles/investigation

- Methodology: https://casrai.org/credit/roles/methodology

- Project administration: https://casrai.org/credit/roles/project-administration

- Resources: https://casrai.org/credit/roles/resources

- Software: https://casrai.org/credit/roles/software

- Supervision: https://casrai.org/credit/roles/supervision

- Validation: https://casrai.org/credit/roles/validation

- Visualization: https://casrai.org/credit/roles/visualization

- Writing — original draft: https://casrai.org/credit/roles/writing-original-draft

- Writing — review & editing: https://casrai.org/credit/roles/writing-review-editing

[Generated with the CASRAI CRediT Statement Builder — https://casrai.org/tools/statement-builder]

## Acknowledgements

We graciously thank the Brun Lab for providing *Henriciella aquimarina* and *Henriciella marina*. *H. mobilis* KCTC 52576 (isolated from the West Pacific Ocean) and *H. pelagia* KCTC 52577 (isolated from the Eastern Pacific Ocean) were obtained from the Korean Collection of Type Cultures (KCTC). *H. marina* JCM 15116 was provided by the RIKEN BRC through the National BioResource Project of the MEXT, Japan. We thank the University of Missouri Electron Microscopy Core (EMC) as well as the Excellence in Electron Microscopy Grant for training and supervision of AMR in electron microscopy. Finally, we are grateful for editorial feedback on drafts from Brody Aubry, Pamela J. Brown, and Breah LaSarre.

